# Neural correlates of winning and losing fights in poison frog tadpoles

**DOI:** 10.1101/2020.01.27.922286

**Authors:** Eva K Fischer, Harmony Alvarez, Katherine M Lagerstrom, Jordan E McKinney, Randi Petrillo, Gwen Ellis, Lauren A O’Connell

## Abstract

Aggressive competition for resources among juveniles is documented in many species, but the neural mechanisms regulating this behavior in young animals are poorly understood. In poison frogs, increased parental care is associated with decreased water volume of tadpole pools, resource limitation, and aggression. Indeed, the tadpoles of many poison frog species will attack, kill, and cannibalize other tadpoles. We examined the neural basis of conspecific aggression in Dyeing poison frog (*Dendrobates tinctorius*) tadpoles by comparing individuals that won aggressive encounters, lost aggressive encounters, or did not engage in a fight. We first compared patterns of generalized neural activity using immunohistochemical detection of phosphorylated ribosomes (pS6) as a proxy for neural activation associated with behavior. We found increased neural activity in the medial pallium and preoptic area of loser tadpoles, suggesting the amphibian homologs of the mammalian hippocampus and preoptic area may facilitate loser-associated behaviors. Nonapeptides (arginine vasotocin and mesotocin) and dopamine have been linked to aggression in other vertebrates and are located in the preoptic area. We next examined neural activity specifically in nonapeptide- and tyrosine-hydroxylase-positive cells using double-label immunohistochemistry. We found increased neural activity specifically in the preoptic area nonapeptide neurons of winners, whereas we found no differences in activity of dopaminergic cells among behavioral groups. Our findings suggest the neural correlates of aggression in poison frog tadpoles are similar to neural mechanisms mediating aggression in adults and juveniles of other vertebrate taxa.

## 1. INTRODUCTION

Aggressive interactions among juveniles are widespread and of ecological and evolutionary importance, despite being less common than among adults. Juveniles are not yet reproductive and rarely hold territories [1–3], and so aggressive interactions arise primarily from competition for nutritional resources that are often provided through parental provisioning. Aggressive interactions among juveniles range from playful, as among litter mates of many mammalian species [4–6], to competitive, as when chicks beg to be fed [7]. Sometimes these interactions can even be lethal, for example when a single dominant nestling kills its siblings [8,9]. When cannibalism occurs, competition and physical aggression can serve to simultaneously defend existing nutritional resources and to provide additional nutrition through consumption of conspecifics [10–12]. While there is ample theory surrounding the evolutionary significance of aggression - especially among siblings - the neural mechanisms underlying juvenile aggression remain poorly understood.

The bulk of studies examining physiological and neural mechanisms of aggression come from adult males, in which sex steroids most often coordinate aggressive behavior [13]. For example, both testosterone and its local conversion to estradiol by aromatase has been linked to aggression in several vertebrate species [14–17]. However, sex steroids are not promising candidates in pre-pubescent, reproductively immature juveniles. What is known about neural mechanisms of aggression in juveniles comes from studies on play fighting in rodents [18–21]. For example, in studies with hamsters and rats, the nonapeptide vasopressin and its signaling through the V1a receptor in the hypothalamus facilitates play fighting in juveniles rodents [18,19]. Additionally, dopaminergic signaling through the D2 receptor facilitates play behavior in juvenile rats [22]. However, play fighting is behaviorally, functionally, and molecularly distinct from violent fighting that can lead to physical damage and death in both adults and juveniles [5]. Given this gap in knowledge of how the brain coordinates of violent aggression in juveniles, we examined neural correlates of aggression in poison frog tadpoles.

Ecological and evolutionary feedback between parental care and offspring aggression influence both juvenile and adult behavior in poison frogs of Central and South America (Family Dendrobatidae). In contrast to most frogs, poison frogs lay their eggs terrestrially and must transport their aquatic larvae to water upon hatching. Species vary in their preferred pool size for tadpole deposition, where smaller pools tend to be predator-free but also nutrient poor, creating a trade-off between safety and nutrition [23]. Increased parental care in dendrobatid poison frogs is associated with resource limitation [23,24] and aggression in tadpoles [5,25,26]. Poison frog tadpoles of multiple species cannibalize con- and hetero-specific eggs and tadpoles [12,25,27,28]. Furthermore, in some poison frog species, tadpole aggressive behavior occurs independent of food limitation [29], suggesting aggression serves to defend access to parental resources, rather than as a means of direct resource acquisition. While other anuran larvae also exhibit cannibalism in response to resource limitation [30–32], an association with parental care is unusual in frogs, and the consequences of tadpole aggression for offspring survival appear to drive complex decisions about tadpole transport and deposition in poison frog parents [12,25,33].

In the present study, we explored the neural correlates of tadpole aggression in a single poison frog species, the Dyeing poison frog (*Dendrobates tinctorius*). We compared patterns of neural activity in eleven candidate brain regions across the brain as well as specifically in neuromodulatory cell types of tadpoles that won aggressive encounters, lost aggressive encounters, or did not engage in a fight. We chose nonapeptidergic neurons (expressing vasotocin and/or mesotocin, the non-mammalian homologues of arginine vasopressin and oxytocin) and dopaminergic neurons as candidates because these neuromodulators have been implicated in aggressive behavior [34,35] including in juveniles [18,20,36]. To our knowledge, this study is the first to examine the neural correlates of violent aggression in amphibians and suggests similar neural mechanisms to those documented in other vertebrate taxa.

## 2. METHODS

### 2.1 Animals

All *Dendrobates tinctorius* tadpoles used for this study were captive bred in our poison frog colony. We established adult breeding pairs following standard procedures in our laboratory. Briefly, one adult male and one adult female were housed together in a 45×45×45 cm terraria containing moss substrate, live plants, a shelter, egg deposition sites, and a water pool for tadpole deposition. Terraria were automatically misted ten times daily, and frogs were fed live *Drosophila* fruit flies dusted with vitamins three times per week. We monitored breeding pairs daily for tadpole transport from the egg deposition site to the water pool. Transported tadpoles were removed from water pools and transferred to individual rearing containers within a larger aquarium where they remained for the duration of development. Water in these large holding aquaria was maintained at 26-28°C and constantly recirculated, such that all tadpoles were exposed to shared water parameters. Tadpoles were fed a diet of brine shrimp flakes and tadpole pellets (Josh’s Frogs, Owosso, MI, USA) three times weekly. In addition, rearing containers contained sphagnum moss and tadpole tea leaves as extra nutrient sources. We checked tadpole health daily and performed a partial water change and water parameter check once weekly. All procedures in this study were approved by the Harvard University Animal Care and Use Committee (Protocol #17-02-293).

### 2.2 Tadpole behavior

Prior to behavioral trials, we randomly sorted tadpoles into size-matched pairs, with all tadpoles used in this study approximately between Gosner stages 30 – 34 (no substantial limb development). We did not exclude sibling pairs as poison frog tadpoles exhibit aggression and cannibalism indiscriminate of kinship [26,27,37]. We size-matched pairs because larger tadpoles are known to win aggressive encounters [26,27,37], and we were interested in neural correlates independent of this physical size advantage. To distinguish individuals during behavioral trials, we stained one randomly chosen tadpole from each pair using neutral red dye (Sigma-Aldrich, St. Louis, MO, USA) following previously established methods [38] and preliminary testing in our lab. Immediately prior to behavioral testing, tadpoles were removed from their rearing containers and placed in 100mL of 0.00125% neutral red in pre-warmed frog holding water for 30 minutes, which turned them pink. This staining is temporary and has no documented impacts on tadpole growth or survival [39,40]. To control for the effects of handling stress, the other member of each pair underwent a sham procedure in which it was similarly removed from its rearing container and placed in 100 mL of pre-warmed holding water for 30 minutes. This staining/sham procedure was immediately followed by behavioral testing.

At the start of each trial, members of a pair were simultaneously placed into a circular arena (5cm diameter, 10cm height) filled with 100 mL of pre-warmed, conditioned frog water. Tadpole pairs were then monitored for aggressive behavior. If no aggression occurred after 60 minutes, trials were stopped and animals returned to their home containers. If aggression occurred, we recorded the latency to first attack and the identity of the attacker. Fights were observed for an additional 45 minutes following this first attack and the ‘winner’ and ‘loser’ identified. This distinction is readily apparent, as winners perform many more aggressive behaviors and losers sustain more tissue damage. We observed only two trials during which we could not identify a clear winner and we excluded these trials from further analysis. Forty-five minutes after fight onset, we terminated trials, took photos of both the winner and loser, euthanized tadpoles with an overdose of benzocaine, and fixed the entire tadpole body in 4% paraformaldehyde. In addition to winners and losers, we collected day- and time-matched control tadpoles from pairs subjected to identical experimental procedures, but who displayed no aggressive interactions. Thus, while pairs differed in total trial duration, the timing of tissue collection following fight onset was consistent across pairs that fought and time-matched in controls. Using our design, roughly half of the dyads fought, facilitating collection of control animals. We kept only one member of each control pair to avoid pseudo-replication. In addition to live observation, all behavioral trials were filmed from above using Samsung HMX-F90 video cameras. Two observers, blind to tadpole identity, quantified behavior from videos, one quantified aggressive behavior using JWatcher software [41] and the other quantified submissive (loser) behavior using BORIS [42] (N=18 per group). For aggressive behavior, we quantified the latency to first attack, the number of attacks, the duration of each attack, the total time attacking, and the average attack duration. For submissive behavior we quantified the number of flights (fleeing events), the duration of each flight, the total time fleeing, and the average flight duration. All behaviors were quantified for both member of each winner/loser pair. We did not analyze behavior for control tadpoles as they, by definition, exhibited no aggressive behavior.

### 2.3 Immunohistochemistry

Whole tadpole bodies were placed into 4% paraformaldehyde at 4°C overnight, rinsed in 1X PBS, and transferred to a 30% sucrose solution for dehydration. Once dehydrated, tadpoles were embedded in mounting media (Tissue-Tek® O.C.T. Compound, Electron Microscopy Sciences, Hatfield, PA, USA), rapidly frozen, and stored at −80°C until cryosectioning. We sectioned the whole tadpole, including the brain, into three coronal series at 10μm. These sections were thaw-mounted onto SuperFrost Plus microscope slides (VWR International, Randor, PA, USA), allowed to dry completely, and then stored at −80°C.

We used an antibody for phosphorylated ribosomes (pS6; phosphor-S6 Ser235/236; Cell Signaling, Danvers, MA, USA) to assess levels of neural activation across the brain. Ribosomes become phosphorylated when neurons are active and pS6 thus serves as a general marker of neural activity, similar to immediate early genes [43]. We followed standard immunohistochemical procedures for 3’,3’-diaminobenzadine (DAB) antibody staining. Briefly, we quenched endogenous peroxidases using a 30% sodium hydroxide solution, blocked slides in 5% normal goat serum to reduce background staining, and incubated slides in primary antibody (rabbit anti-pS6 at 1:500 in blocking solution) overnight. The next day, the slides were washed twice in 1X PBS and then incubated in secondary antibody for 2 hours, followed by avidin-biotin complex (Vectastain Elite ABC Kit; Vector Laboratories, Burlingame, CA, USA) solution incubation for two hours, and treatment with DAB (Vector Laboratories, Burlingame, CA, USA) for 2 min. We rinsed slides with 1X PBS before and after all of the above steps. Finally, slides were rinsed in water, counterstained with cresyl violet, dehydrated in a series of ethanol baths (50%, 75%, 95%, 100%, 100%), and cleared with xylenes prior to cover slipping with permount (Fisher Scientific, Hampton, NH, USA).

Using the two remaining series of sections, we combined the pS6 antibody with a general nonapeptide antibody recognizing both vasotocin and mesotocin or a tyrosine hydroxylase (TH, the rate-limiting step in dopamine synthesis) antibody for standard double-labelled fluorescent immunohistochemistry. Briefly, we blocked slides in 5% normal goat serum to reduce background staining, incubated slides in both primary antibodies (rabbit anti-pS6 (Invitrogen; cat# 44-923G) at 1:500 and anti-vasopressin (PS 45, a generous gift from Hal Grainger) or mouse anti-tyrosine hydroxylase (EMD Millipore; cat# MAB318) at 1:1000) in blocking solution overnight, and incubated slides in a mix of fluorescent secondary antibodies (AlexaFlour 488 anti-rabbit and AlexaFlour 594 anti-mouse at 1:200 in blocking solution) for 2 hours. We rinsed slides with 1X PBS before and after all of the above incubations and rinsed slides in water prior to cover slipping using Vectashield with DAPI (Vector Laboratories, Burlingame, CA, USA). Although we used an anti-vasopressin (PS-45) antibody, immunoblocking the antibody with either vasotocin or mesotocin peptides alone overnight did not block fluorescent signal. However, both peptides combined blocked all signal and thus we refer to these cells as nonapeptide cells generally, rather than vasotocin neurons specifically.

### 2.4 Microscopy & cell counts

Stained brain sections were photographed at 20X on a Leica compound light microscope connected to a QImaging Retiga 2000R camera and a fluorescent light source. We quantified labeled cells from photographs using FIJI image analysis software [44]. Brain regions were identified within and across individuals using a *D. tinctorius* brain atlas constructed in our lab [45]. We measured the area of candidate brain regions and counted all labeled cells within a given region. We quantified cell number in a single hemisphere for each region in each section where that region was visible.

For brightfield pS6 staining (N=10-12 per group), we quantified cell number in the basolateral nucleus of the stria terminalis (BST), the dorsal pallium (Dp), the lateral septum (Ls), the medial pallium (Mp; homolog of the mammalian hippocampus), the anterior preoptic area (aPOA), the medial preoptic area (mPOA), the suprachiasmatic nucleus (SC), the striatum (Str), the posterior tuberculum (TP; homolog of the mammalian ventral tegmental area), the ventral hypothalamus (VH), and the ventral pallium (VP).

For nonapeptide (N=8-9 per group) and TH (N=6 per group) staining, each section was visualized at three fluorescence wavelengths (594nm, 488nm, 358nm) and images were pseudo-colored to reflect these red (pS6), green (nonapeptides or TH), and blue (DAPI) spectra. We used DAPI nuclear staining to identify brain regions. For nonapeptides, we quantified the number of vasotocin and mesotocin positive cells (green), pS6 positive cells (red), and co-labeled cells (yellow) only in the preoptic area and suprachiasmatic nucleus, as this is where nonapeptide cell bodies are found in our study species. As dopamine neurons are much more widespread in the brain, we quantified the number of TH positive cells (green), pS6 positive cells (red), and co-labeled cells (yellow) in all brain regions listed above.

### 2.5 Statistical analyses

All statistical analyses were performed in SAS (SAS 9.4; SAS Institute for Advanced Analytics) and data visualizations in R (version 3.5.0; the R Foundation for Statistical Computing). We began by testing for potential confounding effects in our behavior designs. We tested for differences in body length, body width, and tail length between winner and loser pairs. We included ‘trial’ as a random effect in these analyses to account for the identity of winner and loser pairs. We also tested for behavioral differences between winner and loser pairs for attack number, average attack duration, flight number, and average flight duration. Due to non-normality and unequal variances in the data, we ran these models using a negative binomial distribution.

We tested whether staining color predicted fight outcome, the number of attacks performed, or the number of attacks received. Due to an uneven distribution of pink/control stained winners versus losers, we also tested the effect of color on neural activity in control animals which exhibited no aggressive behavior in order to rule out the likelihood that staining directly influenced neural activity. We ran this later model separately for pS6, TH, and nonapeptide cell counts. pS6 and TH models included cell number, brain region, and their interaction as fixed effects and tadpole identity as a random effect. We did not include brain region as a fixed effect in the nonapeptide analyses because these cell bodies are restricted to the preoptic area and suprachiasmatic nucleus and we combined cell counts from these regions.

We next tested for differences in neural activity based on behavioral group (winner, loser, or control). We ran analyses separately for pS6, TH, and nonapeptide cell counts. pS6 models included cell number, brain region, and their interaction as fixed effects. As we counted TH and co-labeled neuron across multiple brain sections for each individual, we included tadpole identity as a random effect. In addition, we included the log of brain size as a covariate to control for size differences between brain regions and brain size differences among tadpoles. We used Tukey-corrected post hoc analyses to determine which brain regions had significant differences in pS6 positive cell counts.

For TH, we ran models to compare: (1) the total number of TH-positive neurons using a negative binomial distribution appropriate for count data with unequal variances and the log of the total number of brain sections quantified as an offset variable; and (2) the proportion of active TH neurons (i.e. the total number of neurons co-labeled for pS6 and TH divided by the total number of TH positive neurons) using a binomial distribution. We included behavioral group, brain region, and their interaction as fixed effects in both models. For the nonapeptides, we ran analyses similar to those for TH, but without brain region as a fixed effect. We tested (1) the total number of nonapeptide-positive neurons using a negative binomial distribution and the log of the total number of brain sections quantified as an offset variable; and (2) the proportion of active nonapeptide neurons (i.e. the total number of neurons co-labeled for pS6 and nonapeptides divided by the total number of nonapeptide positive neurons) using a binomial distribution. We included behavioral group as a fixed effect. For both TH and nonapeptide analyses we included the log of the number of sections counted as an offset variable to control for differences in sampling (e.g. due to issue damage and/or differences in tadpole body size) and again used Tukey-correction for multiple hypothesis testing to adjust p-values for all post hoc comparisons.

Finally, we tested for associations between neural activity and behavior. We excluded control animals from these analyses as they by definition exhibited no aggressive behavior. We first tested for differences between winners and losers in the total number of attacks, total number of flights, average attack duration, and average flight duration. We next tested for linear relationships between neural activity and each behavioral metric. We ran separate models for total nonapeptide neuron number and active nonapeptide neuron number. We included behavioral group, neuron number, and their interaction as fixed effects predicting behavior (attack number, attack duration, flight number, flight duration), and the log number of sections quantified as an offset variable. A significant main effect of neuron number indicates a group-independent relationship between neural activity and behavior. In contrast, a significant interaction indicates that the relationship between neural activity and behavior differs by behavioral group. We did not evaluate group differences from these models as we explicitly tested for them above and included behavioral group here only to control for group differences in neuron number.

## 3 RESULTS

### 3.1 Behavioral differences

Winners attacked on average 30 times (se ±4.7) with an average attack duration of 0.92 seconds, significantly more than losers who attacked on average 5 times (se ±1.5) with an average attack duration of 0.19 seconds (Fig. 1a; F_1,34_=27.22, p<0.0001). Conversely, losers fled on average 96 times (se ±11.5), significantly more than winners who fled on average 7 times (se ±0.9) (Fig. 1b; F_1,34_=194.25, p<0.0001). There was a trend toward longer average attack duration in winners (Fig. 1c; F_1,34_=4.02, p=0.0530), but no difference in average flight duration between winners and losers (Fig. 1d; F_1,34_=1.52, p=0.2263). Controls by definition performed zero attacks. There were no differences in body length, body width, or tail length between winner and loser pairs.

**Figure 1.**
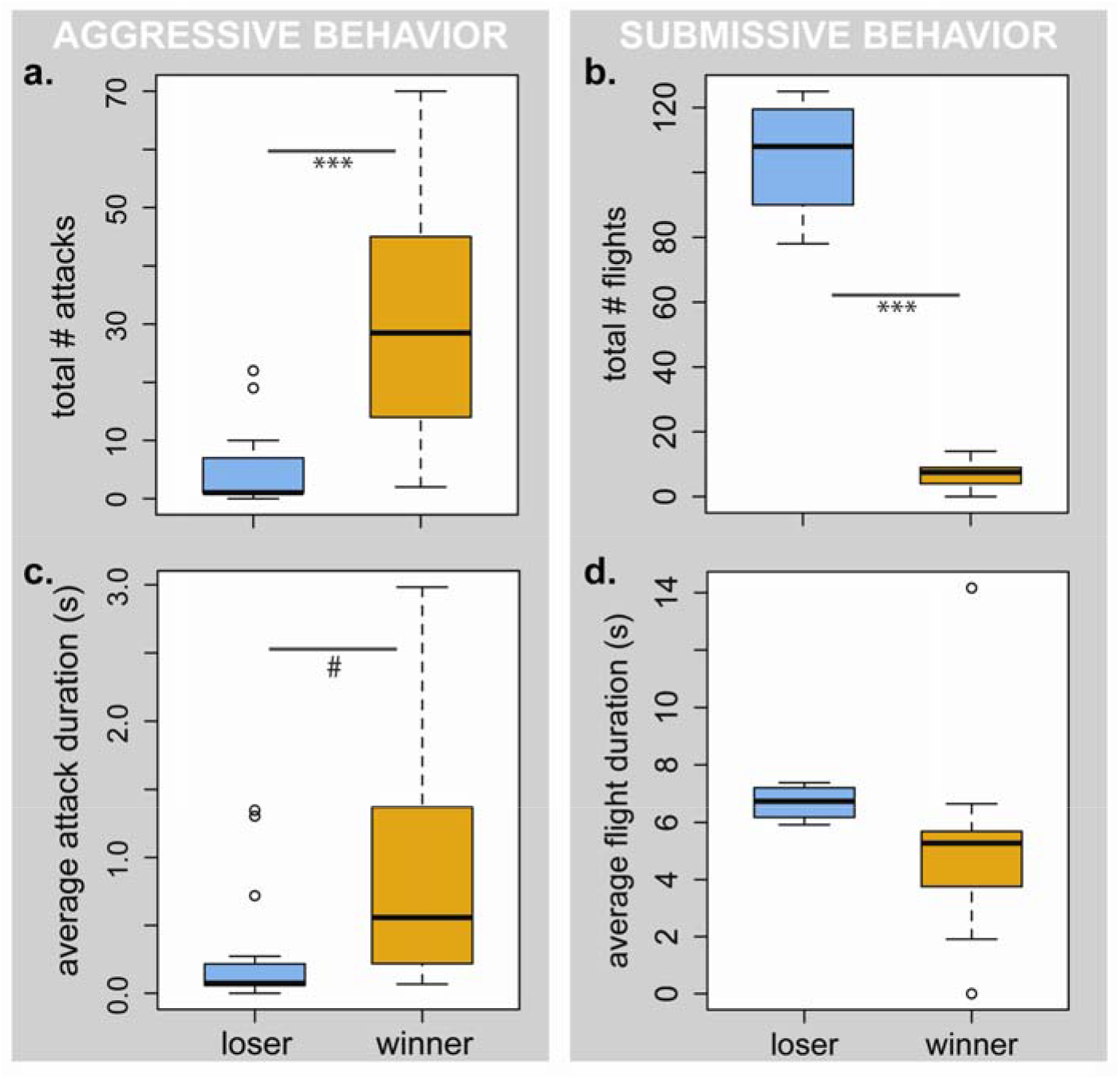
Winning and losing behavior. Winners (a) attacked significantly more and (b) fled significantly less than losers. (c) Average attack duration was marginally longer in winners, but (d) there was no difference in average flight duration between winners and losers (N=18 per behavioral group). Boxplots show median (black bar), the first and third quartiles (box edges), 1.5 times the interquartile range (whiskers), and any outliers (dots). # p<0.1, *** p<0.001.

Color treatment did not predict behavioral group (winner, loser, or control; χ^2^=2.29, p=0.130). Nor did color treatment predict attack number (F_1,38_=1.77, p=0.198), average attack duration (F_1,38_=0.27, p=0.607), or the number of attacks received (F_1,38_=1.77, p=0.198). Nonetheless, we noticed a greater proportion of pink tadpoles among the losers (Fig. S1a; χ^2^=6.59, p=0.010). To ensure that this effect of color treatment was not directly influencing neural activity, we compared neural activity between pink and neutral controls (i.e. fight behavior-independent levels of neural activity) and found no significant differences in these analyses (Fig. S1b).

### 3.2 Behavioral group differences in neural activity

Differences in general neural activity depended on behavioral group and brain region (group*region: F_22,3681_=4.41, p<0.001). Post hoc analyses revealed significant group differences in the medial pallium (non-mammalian homolog of the hippocampus; F_22,3681_=4.46, p=0.011) and the anterior preoptic area (F_2,3681_=3.68, p=0.025) (Fig. 2). In the medial pallium, losers had significantly more activity than controls, but losers and winners did not differ, nor did winners and controls. In the preoptic area, losers had significantly more activity than winners, but neither differed from controls. Results for all brain regions are shown in Table 1 and Figure S2.

**Table 1.** Post hoc test of regional differences in neural activity among behavior groups. Abbreviations: BST = basolateral nucleus of the stria terminalis; Dp = dorsal pallium; LS = lateral septum; Mp = medial pallium; aPOA = anterior preoptic area; mPOA = medial preoptic area; SC = suprachiasmatic nucleus; Str = striatum; TP = posterior tuberculum; VH = ventral hypothalamus; VP = ventral pallium.

| brain region | df num | df den | F | p |
| --- | --- | --- | --- | --- |
| BST | 2 | 3282 | 0.58 | 0.559 |
| Dp | 2 | 3282 | 0.98 | 0.374 |
| Ls | 2 | 3282 | 0.64 | 0.526 |
| Mp | 2 | 3282 | 4.46 | **0.011** |
| aPOA | 2 | 3282 | 3.68 | **0.025** |
| mPOA | 2 | 3282 | 1.37 | 0.254 |
| SC | 2 | 3282 | 1.33 | 0.266 |
| Str | 2 | 3282 | 1.03 | 0.358 |
| TP | 2 | 3282 | 0.54 | 0.585 |
| VH | 2 | 3282 | 0.31 | 0.734 |
| VP | 2 | 3282 | 2.13 | 0.119 |

**Figure 2.**
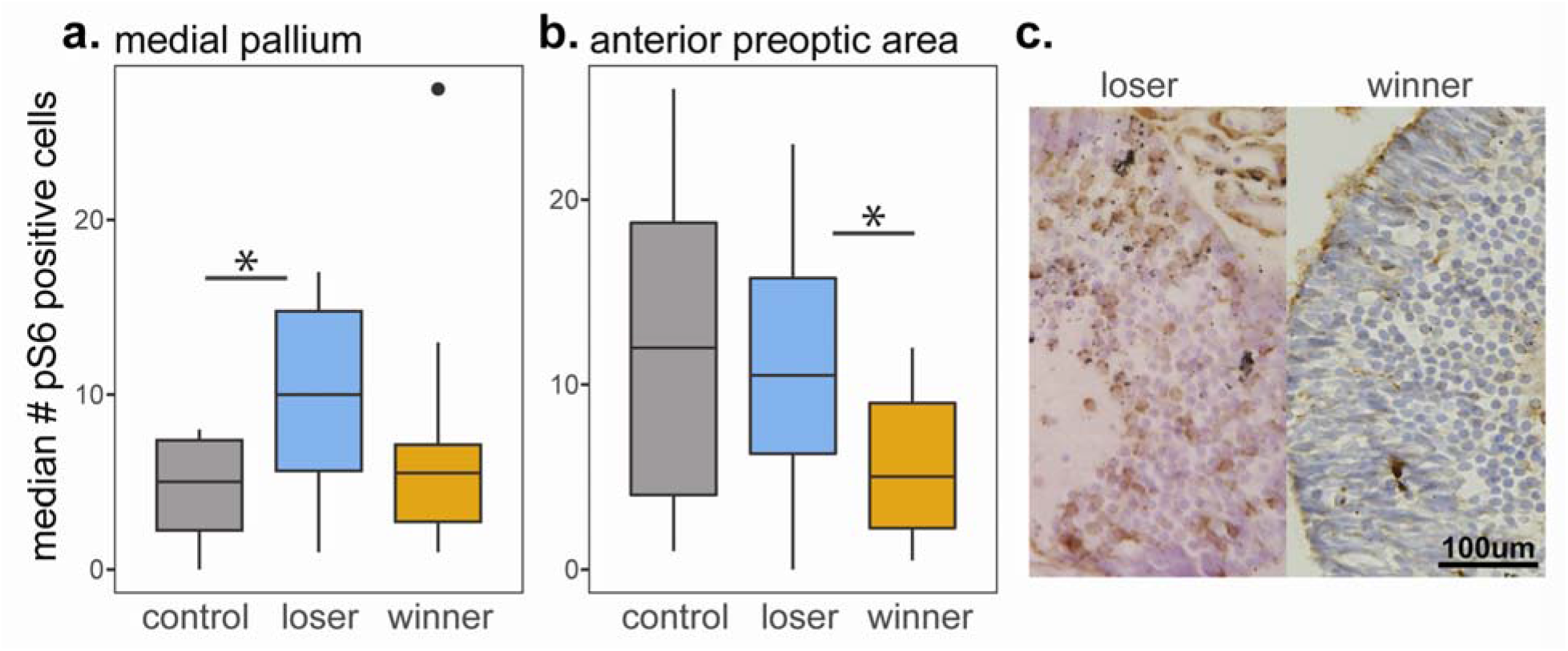
Differences in neural activity based on behavior. We quantified neural activity using a pS6 antibody for phosphorylated ribosomes. We found differences in the number of pS6 positive cells among behavioral groups in (a) the medial pallium and (b) the anterior preoptic area. In the medial pallium, losers had significantly more activity than controls and marginally more activity than winners, while winners and controls did not differ. In the anterior preoptic area, losers had higher activity than winners, but neither differed from controls. Representative staining from the preoptic area is shown in (c). Results for all brain regions are in Table 1 and Figure S3. N=11 winners, N=12 losers, and N=10 controls. Boxplots show median (black bar), the first and third quartiles (box edges), 1.5 times the interquartile range (whiskers), and any outliers (dots).

In addition to patterns of generalized neural activity, we quantified nonapeptide (vasotocin and mesotocin, amphibian homologs of the mammalian arginine vasopressin and oxytocin peptides, respectively) and tyrosine hydroxylase (TH, the rate-limiting step in dopamine synthesis) positive neurons. While we found no differences in the total number of nonapeptide positive cells (Fig. 3; group: F_2,27_=0.36, p=0.699), we observed a difference in the proportion of active nonapeptide cells between behavioral groups (Fig. 3; group: F_2,27_=11.00, p=0.0004). Winners had a greater proportion of active nonapeptide neurons than controls (t_23_=−4.66, p=0.0003) and a marginally greater proportion of active nonapeptide neurons than losers (t_23_=−2.30, p=0.075), who also had a marginally greater proportion of active nonapeptide neurons than controls (t_23_=−2.29, p=0.075). We found no overall group or brain region specific differences in the total number of TH positive cells (Fig. S3) or the proportion of active TH cells (Fig. S4).

**Figure 3.**
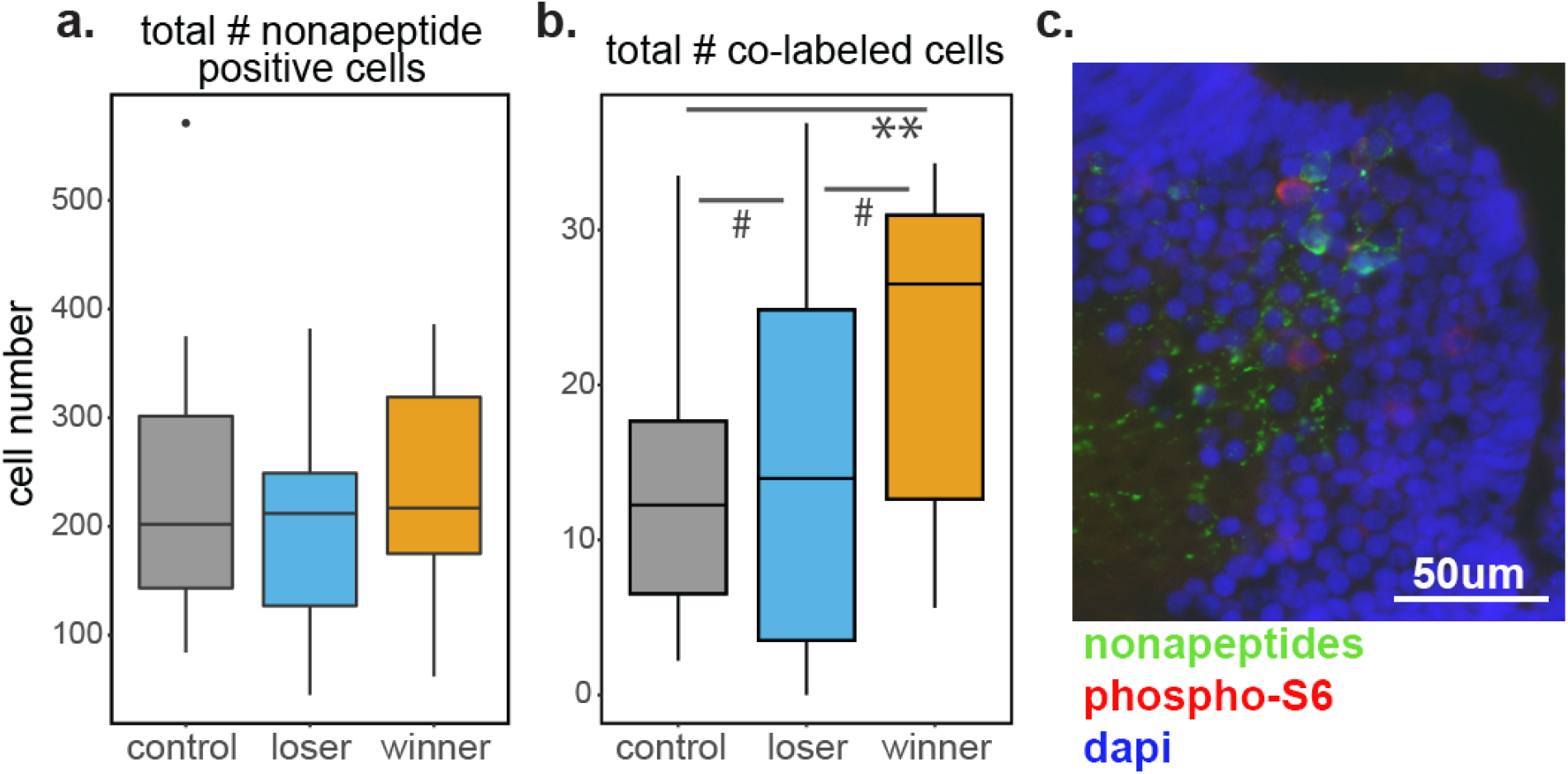
Nonapeptides and aggressive behavior. (a) No differences among behavioral groups in the total number of nonapeptide positive neurons, but (b) winners have a greater number of active nonapeptidergic neurons. (c) Representative micrograph showing nonapeptide positive cells (green), pS6 positive cells (red), co-labeled cells (yellow), and dapi nuclear staining (blue). N=9 winners, N=9 losers, and N=8 controls. Boxplots show median (black bar), the first and third quartiles (box edges), 1.5 times the interquartile range (whiskers), and any outliers (dots). # p<0.1, ** p<0.01.

### 3.3 Associations between neural activity and behavior

In light of within group variation in how much individuals fought before winner/loser identity was established, we explored relationships between neural activity and behavior. We did not include control animals in these analyses as they by definition did not engage in any aggressive behavior. We found a significant relationship between nonapeptidergic activity and aggressive, but not submissive, behavior. The total number and the active number of nonapeptidergic neurons did not have a significant linear relationship with attack number. However, the total number of nonapeptide neurons (F_1,14_=18.11, p=0.0008) and the active number of nonapeptide neurons (F_1,14_=40.18, p<0.0001) predicted attack duration independent of behavioral group (Fig. 4). There was no relationship between nonapeptide neuron number or activity with either flight number or duration.

**Figure 4.**
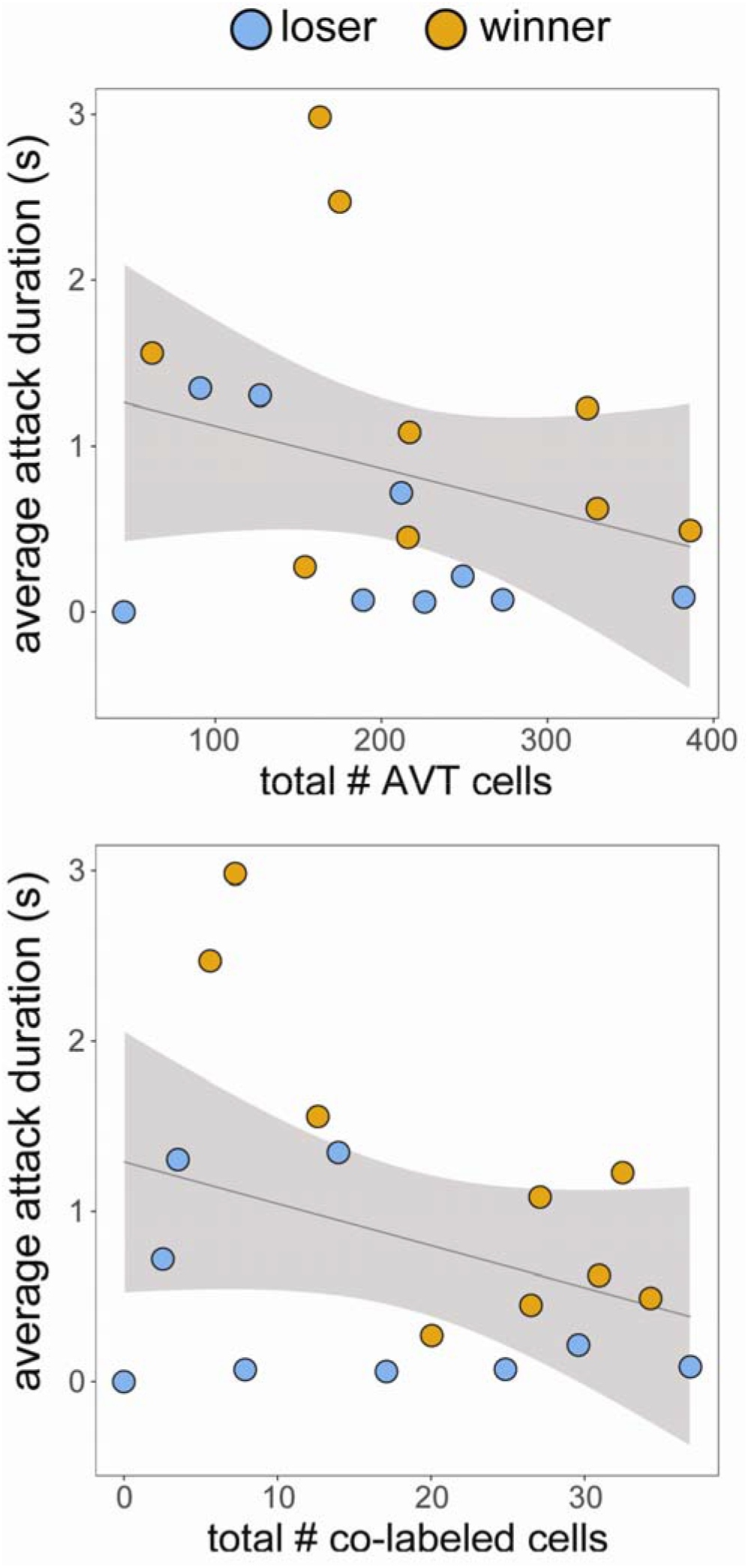
Neural activity and behavior. The total number and total number of active (i.e. co-labeled) nonapeptidergic neurons predicts attack duration in a group-independent fashion.

## 4 DISCUSSION

We examined the neural basis of conspecific aggression in *Dendrobates tinctorius* poison frog tadpoles by comparing patterns of neural activity across eleven brain regions, as well as specifically in nonapetidergic and dopaminergic neurons in tadpoles that won or lost aggressive encounters or did not engage in a fight. To our knowledge, this is the first exploration of the neural correlates of violent juvenile aggression in non-mammals.

### 4.1 General brain activity associated with fight outcome

Comparing patterns of neural activity in winner, loser, and control tadpoles, we found differences in the anterior preoptic area and medial pallium (non-mammalian homolog of the hippocampus). The hippocampus and its non-mammalian homologs are classically implicated in memory, and specifically spatial memory [46–48]. In *D. tinctorius* tadpoles, losers had significantly higher medial pallial activity than controls and marginally higher medial pallial activity than winners. Given the deadly consequences of aggressive encounters between tadpoles, increased medial pallial activity in losers may help tadpoles to avoid the sites of these encounters in future. While such an avoidance strategy is not possible in species that use the smallest tadpole deposition pools, *D. tinctorius* deposit their tadpoles in pools large enough to make avoidance feasible. The neural architecture of the medial pallium is not well understood, and more research should be done to determine which cells are active in loser tadpoles and the role of the neurons in governing behavior during or after aggressive encounters.

The preoptic area plays functionally conserved roles in the regulation of social behavior across vertebrates [49], and we found the anterior preoptic area had increased neural activity in losers compared to winners. In the context of aggression, electrical stimulation of the preoptic area increases aggression in adult rodents [50], birds [51], reptiles [52], and fish [53]. While we found differences in preoptic area activity associated with aggression, we found increased preoptic area activity in losers as compared to winners, the opposite of the relationship suggested by electrical simulation studies. We provide a number of potential explanations for this finding. First, the role of the preoptic area specifically in juvenile aggression remains unresolved. While studies in adults implicate the preoptic area in aggressive behavior, preoptic area lesions in juvenile rats [54] and dogs [55] do not disrupt the performance of or development of aggressive behavior. Second, given the wide-ranging social behavioral functions of the preoptic area, associations between behavior and overall region activity may paint a distinct picture from associations between behavior and the activity of specific neuronal types. We note that controls had intermediate levels of activity that did not differ significantly from either winners or losers. As controls were also exposed to, but did not fight with, conspecifics, this pattern further suggests specificity in the relationship between neural activity and distinct types of social interactions. Our exploration of associations between the activity of two candidate cell types and behavior (see below) suggest that future studies explicitly distinguishing overall neural activity from that of specific cell types will be fruitful.

### 4.2 Neural activity in nonapeptide and dopaminergic cells

Given behavioral and neural activity differences between winners and losers, we next asked whether nonapeptides and dopamine known to mediate aggression in other species were associated with fight outcome. While we found overall increased preoptic area activity in losers, we observed increased activity specifically in nonapeptide neurons in winners. This finding is in line with observations in other vertebrates that link nonapeptides and aggressive behavior. Vasopressin/vasotocin have been linked to aggression generally, particularly in males [34]. Oxytocin is most prominently associated with maternal aggression [56], but increased oxytocin neuron activity has also been linked with aggression more generally [57]. Importantly, while this broad association between nonpeptides and aggression is well-established, the precise nature of these relationships differs based on sex, species, and experience [35,58].

In juveniles specifically, changes in aggressive behavior following early life stress are associated with developmental differences in oxytocin and vasopressin systems in rats [19,59]. Increased arginine vasopressin is linked to increased play fighting in juvenile hamsters [18,19], while decreased vasopressin signaling in juvenile *Peromyscus* mice is associated with decreased aggression [60]. Notably, although patterns of arginine vasopressin innervation differ between sexes in adults of many species, this is not the case in juvenile rats [20,60]. It is unknown whether this sex-independent innervation pattern in juveniles is wide-spread. There is no genetic sex marker for poison frogs and thus whether aggression or neural architecture differs by sex is unknown. However, we suggest that nonapeptide signaling provides a promising mechanism for investigating violent juvenile aggression in this and other species.

In contrast to activity differences in nonapeptide neurons among behavioral groups, the total number of nonapeptide neurons did not differ between winners, losers, and controls. We interpret this observation as evidence that fight outcome is associated with the activity of nonapeptide neurons, rather than baseline differences in neuron number. Links between aggressive behavior and neuronal activity, rather than neuron number, make sense in light of the need for animals to integrate internal (e.g. hunger levels, body condition) and external (e.g. size and identity of opponent) cues in order to make appropriate real-time behavioral decisions during aggressive encounters. While all animals used in this study were fight-naïve, social interactions are known to modulate short- and long-term nonapeptide signaling [34,58,61] providing avenues for future investigation.

Because tadpole pairs varied in how much they fought before winner/loser identity was established, we examined relationships between nonapeptidergic activity and behavior in a group-dependent and group-independent manner. We found a group-intendent association between neuropeptidergic activity and aggressive, but not submissive, behavior. We suggest that these observations imply that distinct neuromodulators mediate opposing aggressive and submissive behaviors, and that nonapeptide activity is linked to the performance of aggressive behavior, regardless of fight outcome. Although we cannot presently distinguish neural activity differences associated with differences in fight progression, future studies exploring individual variation in behavior, linking the activity of additional cell types to aggressive and submissive behavior, and disentangling the role of vasotocin versus mesotocin will shed light on these open questions.

In addition to nonapeptides, we examined tyrosine hydroxylase-positive neurons as a proxy for dopamine, which has been linked to differences in aggression among individuals, including specifically in the context of winner/loser effects [62]. We found no differences in either the total number of dopamine neurons or the activity of these neurons across winners, losers, and controls. We emphasize that this does not rule out a role for dopamine in aggression. First, dopamine signaling relies on a number of different receptors whose behavioral roles differ by receptor subtype [62–64], and we did not examine receptor levels. Second, the influence of dopamine signaling on aggressive behavior can vary by species based on the nature of aggressive interactions. For example, previous studies have found that the relationship between aggression and dopamine is mediated by activity differences among aggressive and non-aggressive individuals [61], and that dopamine is associated with willingness to engage in social interactions rather than with aggression directly [36]. Finally, given previous studies demonstrating the role of dopamine in mediating winner/loser effects [62], differences in dopamine signaling may be apparent at timescales outside of the narrow window which we sampled, and/or become apparent only after repeated fight experience. In sum, further studies examining the role of dopamine specifically in the context of winner/loser effects may be valuable.

### 4.3 Experimental limitations

Neutral red and other vital dyes have been used to distinguish individual tadpoles in a variety of experiments with no apparent consequences for growth and survival [39,40]. Furthermore, Wilcox and Lapping (2013) used a behavioral procedure very similar to ours and found no effect of neutral red on fight outcome in a closely related poison frog species, *Dendrobates auratus*. Using the same concentration and timing as these previous studies, we observed an association between neutral red treatment and behavior, where neutral red treated tadpoles were more likely to lose aggressive encounters. This effect did not appear to be mediated by conspecific responses to pink tadpoles, as dye treatment did not predict the number of attacks received. Nor did dye treatment appear to influence fight outcome by directly altering neural activity, as there were no differences in neural activity between neutral red and sham treated tadpoles that did not fight. In other words, differences in neural activity between winners and losers are fight-behavior specific. For this reason, we interpret neural differences in the context of fight outcome, noting that our finding of an association between dye color and fight outcome may be mediated by differences among tadpoles in an unmeasured variable and/or coincidence. In either case, we caution against the existing assumption that this common staining procedure does not influence tadpole behavior.

## 5 Conclusions

We found differences in overall neural activity and specifically in the activation of nonapeptide neurons between tadpoles that won, lost, or did not engage in a fight. While overall neural activity in the preoptic was greater in losers, winners has increased activation specifically in preoptic area nonapeptide neurons. These patterns suggest that – especially for integrative multi-functional nodes of the social decision-making network, such as the preoptic area – differences in overall activity versus the activity of specific neuronal types may have distinct associations with behavior. To our knowledge, this is the first exploration of the neural mechanisms of juvenile aggression in an amphibian. Our findings provide a starting point for research on the neural mechanisms of violent aggression in juvenile amphibians and other vertebrates.

## ACKNOWLEDGEMENTS

We thank Kim L Hoke and Kate L Laskowski for fruitful discussions of statistical analyses, high school teachers Barbara Dorritie (Cambridge Rindge and Latin) and Tammy Fay (Masconomet) for their co-mentorship of high school students (RP and GE, respectively) during their research internships, and all the frog caretakers that maintain our poison frog colony. We thank Julie Butler and two anonymous reviewers for comments on previous versions of the manuscript.

## FUNDING

This work was supported by a Bauer Fellowship from Harvard University and the Rita Allen Foundation to LAO, a National Science Foundation Postdoctoral Fellowship in Biology (NSF-1608997) to EKF, and a Stanford University Biology Summer Undergraduate Research Program Fellowship to HA.

## Notes

### Competing Interest Statement

The authors have declared no competing interest.

