## Supplemental Materials for "Neural correlates of winning and losing fights in poison frog tadpoles"

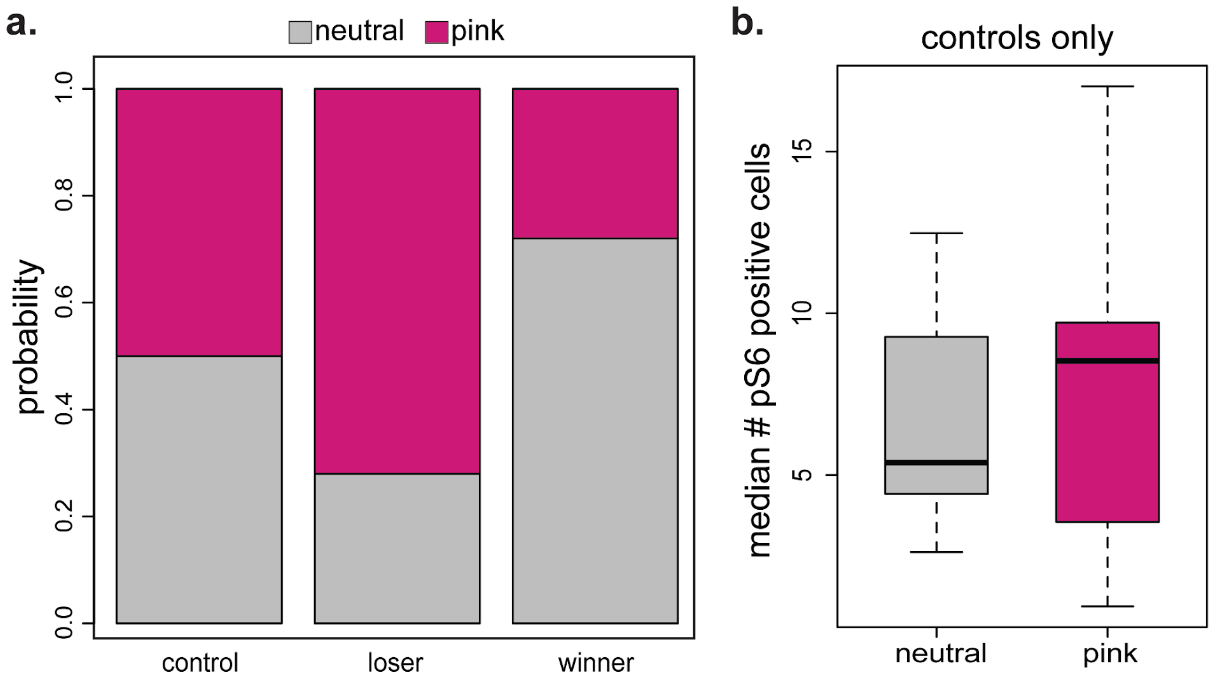


**Figure S1.** Effect of color treatment on fight outcome but not neural activity. (a) Among the tadpoles that fought, those randomly assigned to the pink color treatment (i.e. staining with neutral red dye) were more likely to lose aggressive encounters. (b) No differences in neural activity based on color treatment. Only control tadpoles, in which neural activity is independent of fight outcome, are shown.

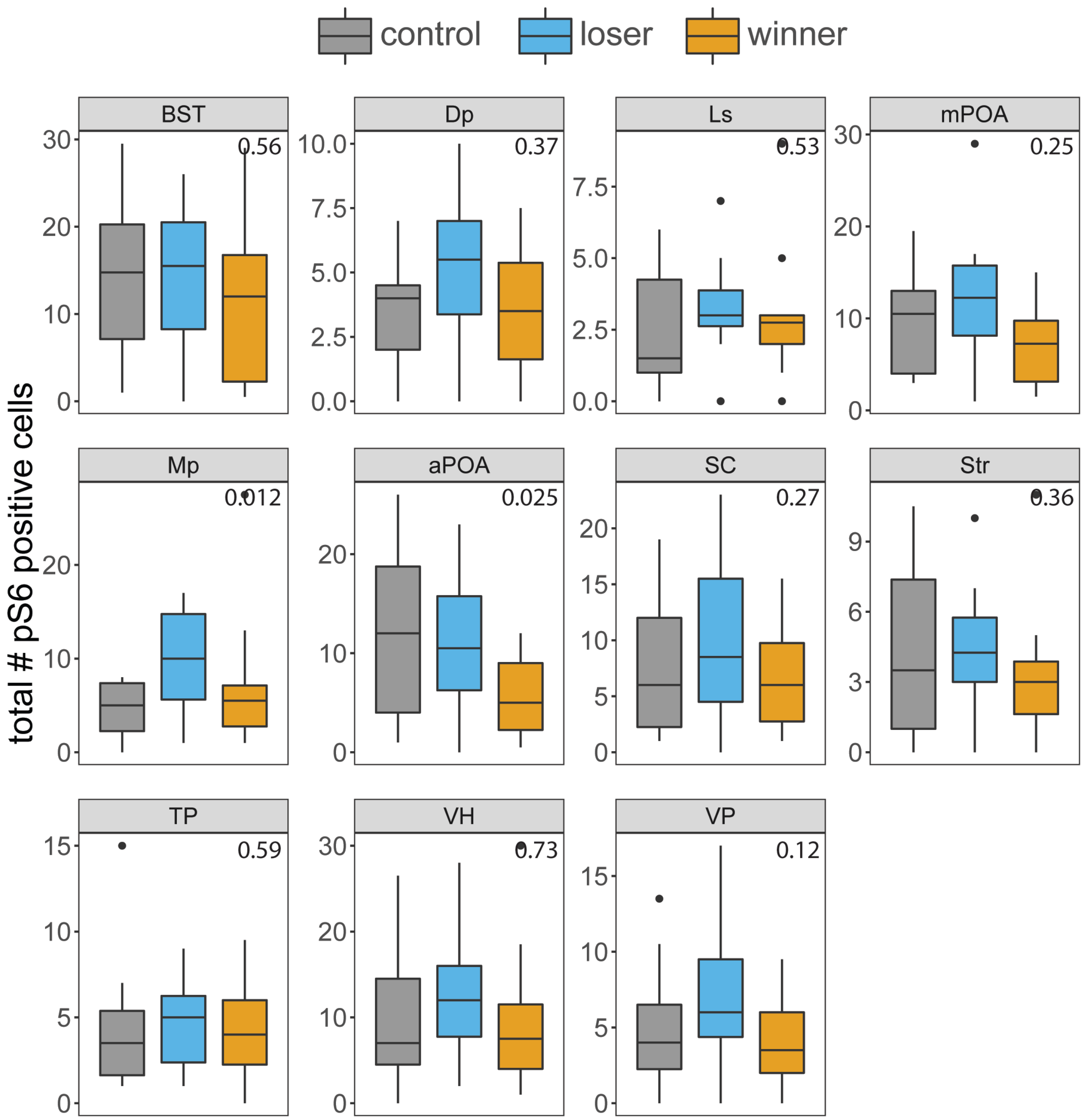


**Figure S2.** Patterns of neural activity among behavioral groups by brain region. Tukey corrected p-values for posthoc comparisons of group differences by brain region are in the top right corner of each plot. We found significant differences in neural activity based on behavioral group in the anterior preoptic area and medial pallium. BST = basolateral nucleus of the stria terminalis; Dp = dorsal pallium; Ls = lateral septum; Mp = medial pallium (homolog of the mammalian hippocampus); aPOA = anterior preoptic area; mPOA = medial; SC = suprachiasmatic nucleus; Str = striatum; TP = posterior tuberculum (homolog of the mammalian ventral tegmental area); VH = ventral hypothalamus; VP = ventral pallium. N=9 winners, N=9 losers, and N=8 controls. Boxplots show median (black bar), the first and third quartiles (box edges), 1.5 times the interquartile range (whiskers), and any outliers (dots).

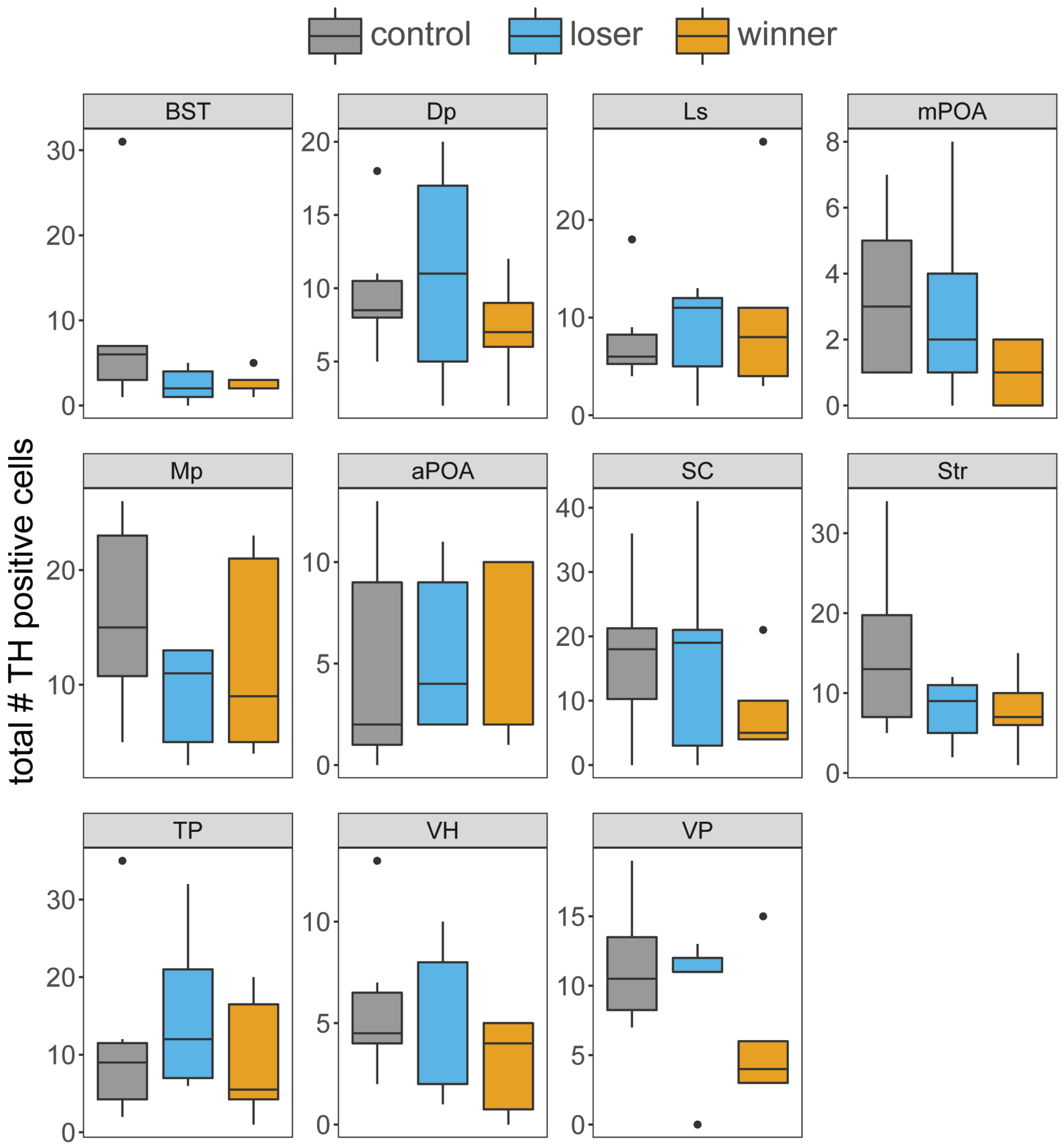


**Figure S3.** No main effect of behavioral group or brain region specific effects of behavioral group on the number of TH positive neurons. BST = basolateral nucleus of the stria terminalis; Dp = dorsal pallium; Ls = lateral septum; Mp = medial pallium (homolog of the mammalian hippocampus); aPOA = anterior preoptic area; mPOA = medial; SC = suprachiasmatic nucleus; Str = striatum; TP = posterior tuberculum (homolog of the mammalian ventral tegmental area); VH = ventral hypothalamus; VP = ventral pallium. N=6 winners, N=5 losers, and N=6 controls. Boxplots show median (black bar), the first and third quartiles (box edges), 1.5 times the interquartile range (whiskers), and any outliers (dots).

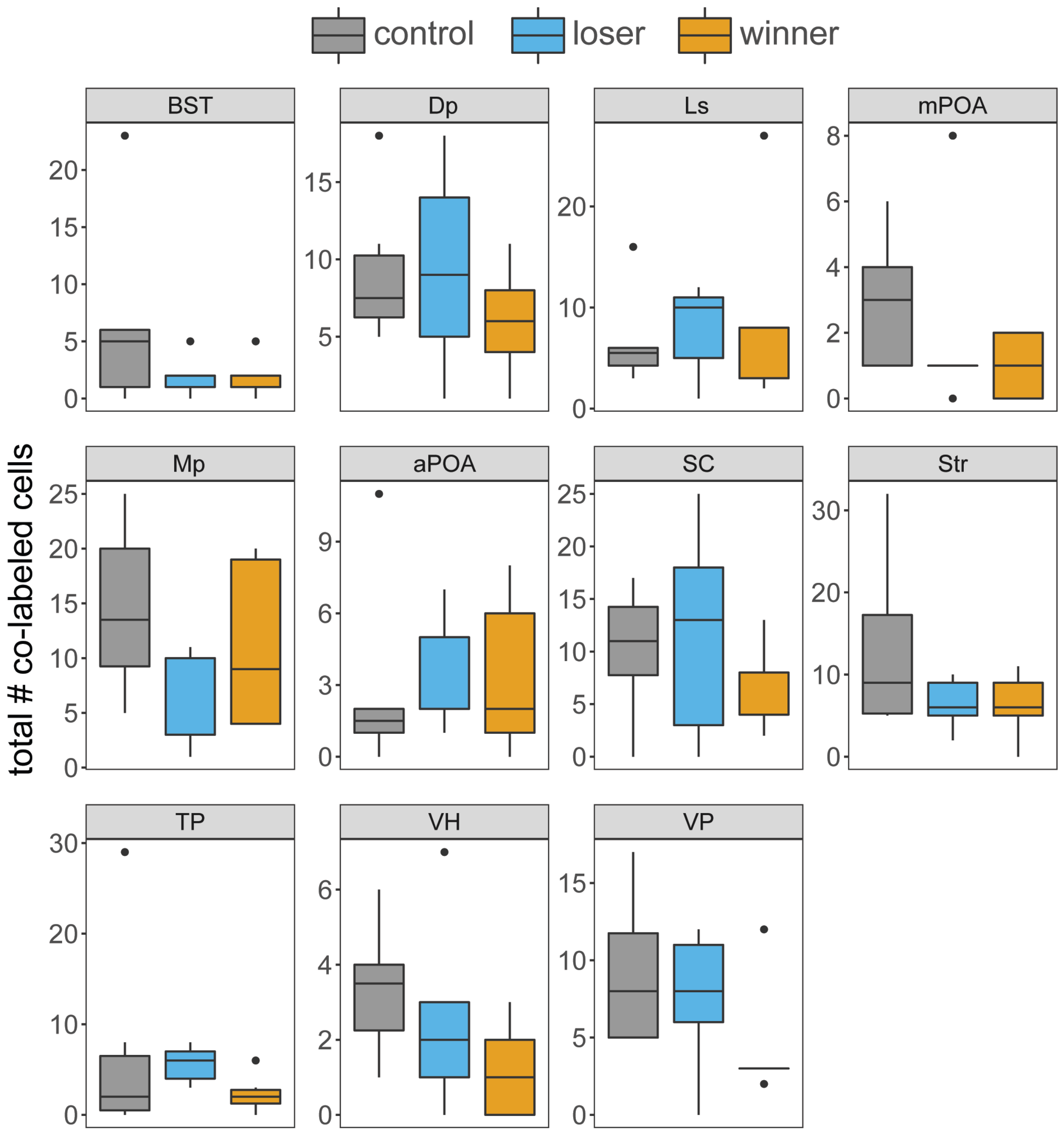


**Figure S4.** No main effect of behavioral group or brain region specific effects of behavioral group in the proportion of active TH neurons. BST = basolateral nucleus of the stria terminalis; Dp = dorsal pallium; Ls = lateral septum; Mp = medial pallium (homolog of the mammalian hippocampus); aPOA = anterior preoptic area; mPOA = medial; SC = suprachiasmatic nucleus; Str = striatum; TP = posterior tuberculum (homolog of the mammalian ventral tegmental area); VH = ventral hypothalamus; VP = ventral pallium. N=6 winners, N=5 losers, and N=6 controls. Boxplots show median (black bar), the first and third quartiles (box edges), 1.5 times the interquartile range (whiskers), and any outliers (dots).
